# Microfluidic Assembly of Synthetic Tissues from Synthetic Cells

**DOI:** 10.64898/2026.11.10.765587

**Authors:** Indraja Sundara Raju, Mostafa Bakouei, Benny Ryplida, Tatiana Avsievich, Caglar Elbuken

## Abstract

Biological tissues demonstrate emergent properties arising from the collective interactions of individual cells. While developmental biology has revealed the molecular mechanisms that control processes such as cell-cell adhesion, cellular rearrangements, and mechanosignalling, understanding how the local interactions lead to tissue scale mechanics remains challenging due to the biochemical complexity of living systems. Minimal synthetic tissue models allow us to isolate and examine the underlying physical principles. Liposomes as synthetic cells provide a biomimetic platform with tunable membrane characteristics and the ability to assemble into more complex structures. We present minimal tissue mimetic models based on synthetic cells that enable quantification to link microscale interactions to macroscale features in minimal synthetic tissues. To facilitate these studies, we have optimized the solvent mediated dewetting method to produce some of the highest throughput. This high yield enabled us to develop tissue mimetic structures in two complementary configurations: a freestanding monolayer and a confined microfluidic assembly. Morphological analysis in freestanding model revealed that adhesion-mediated packing induced significant deformation of synthetic cells, accompanied by changes in circularity and eccentricity. The fluorescent assay in confined model demonstrated that adhesive tissue formation reduced transport rates by decreasing pore size and increasing pathway tortuosity while maintaining mechanical stability under flow. Together, these minimal tissue-mimetic platforms provide a quantitative framework for linking synthetic-cell properties, collective organization, and emergent tissue-scale phenomena.

## 1. Introduction

Bottom-up synthetic biology seeks to answer a fundamental question: how do simple components self-organize to create the complex mechanics of living systems? If we cannot build a living system from its basic components, our understanding of how it works remains incomplete. Synthetic cells provide the simplest platform to reconstruct these life-like functions from non-living materials. The introduction of liposomes as synthetic cells has ushered in a new era of biomimetics and bottom-up synthetic biology. Since their discovery in the early 1960s, remarkable progress has made in engineering synthetic cells.^1–6^ The focus has now shifted from individual synthetic cells to synthetic multicellular systems and origin-of-life investigations. Research is increasingly focused on synthetic tissue construction from these synthetic cells, aiming to exploit their dynamic behaviors and mechanisms. Yet, prototissues, the assembly of organized synthetic cells that can exhibit collective behaviors, are still in their infancy.^7^

Biological tissues demonstrate emergent properties arising from the collective interactions of individual cells rather than the isolated cell characteristics.^8^ Cell-cell adhesion, cellular rearrangements and mechanosignalling ^9^ are ways in which the cells self-organize and collectively respond to external physical and chemical stimuli. While developmental biology has revealed molecular mechanisms controlling these interactions, understanding how they lead to tissue scale mechanics remains challenging due to the biochemical complexity of living systems. ^7^

Research efforts are directed to mimic them in minimal synthetic tissue models as studying these interactions in living tissues is complicated with the numerous signalling pathways and cellular processes.^10–12^ Despite the advances, the invitro minimal models and the quantitative framework connecting microscale properties, spatial organisation and macroscale tissue properties in the synthetic tissue models remain lacking. ^13^

In this study, we address the lack of invitro minimal tissue models required to study the collective behaviour and emergent tissue properties. We develop synthetic tissue-mimetic assemblies from adhesive synthetic cells and investigate their structural and functional properties in both freestanding and confined systems. In the freestanding tissue mimetic model, we investigate the quantification of cell shapes to facilitate the understanding of impact of shape morphology on tissue functions. The confined tissue mimetic model is the platform we have conceptualised to test the transport of bio analytes in the packed assemblies.

## 2. Results and Discussion

### 2.1. Microfluidic production of synthetic cells

In this study we developed tissue mimics by assembling synthetic cells. Giant unilamellar vesicles (GUV) made using the double emulsion template method were the building blocks of our synthetic tissue model. Double emulsions (DE) are converted to liposomes using the solvent – mediated dewetting. DEs were prepared using the hybrid chip^14^ technology discussed in our earlier study (Fig 1b). DEs from the hybrid chip were collected in a well plate (Fig 1a) and incubated overnight at room temperature. DE shells comprised of a lipid dissolvable solvent (hexanol) and an incompatible solvent for lipid dissolution (paraffin oil). On removal of hexanol the DE transition to the intermediary state shown as dewetting synthetic cells in Figure 1c.^15^ Due to the density difference between the inner aqueous phase and outer aqueous phase synthetic cells tend to sediment. The oil droplet floats to the top and the synthetic cell sediments at the bottom of the well plate. Synthetic cell yield in the well plate was as seen in Figure 1d. Then the synthetic cells were transferred to another well plate to minimise impurities. We have observed that leaving the sample in the collected well plate containing the residual oil droplets resulted in formation of aggregates or fibrous structures that contaminated the sample. When sample is transferred to the new well plate, minimal contamination and improved stability was achieved. Moreover, the sample can be stored at 4°C and used for at least two weeks. The high yield and stability achieved with our technique provides robust foundation for the assembly of synthetic tissue model. Adding intercellular adhesion functionality to our synthetic cells we developed two different platforms to study tissue scale properties.

**Figure 1:**
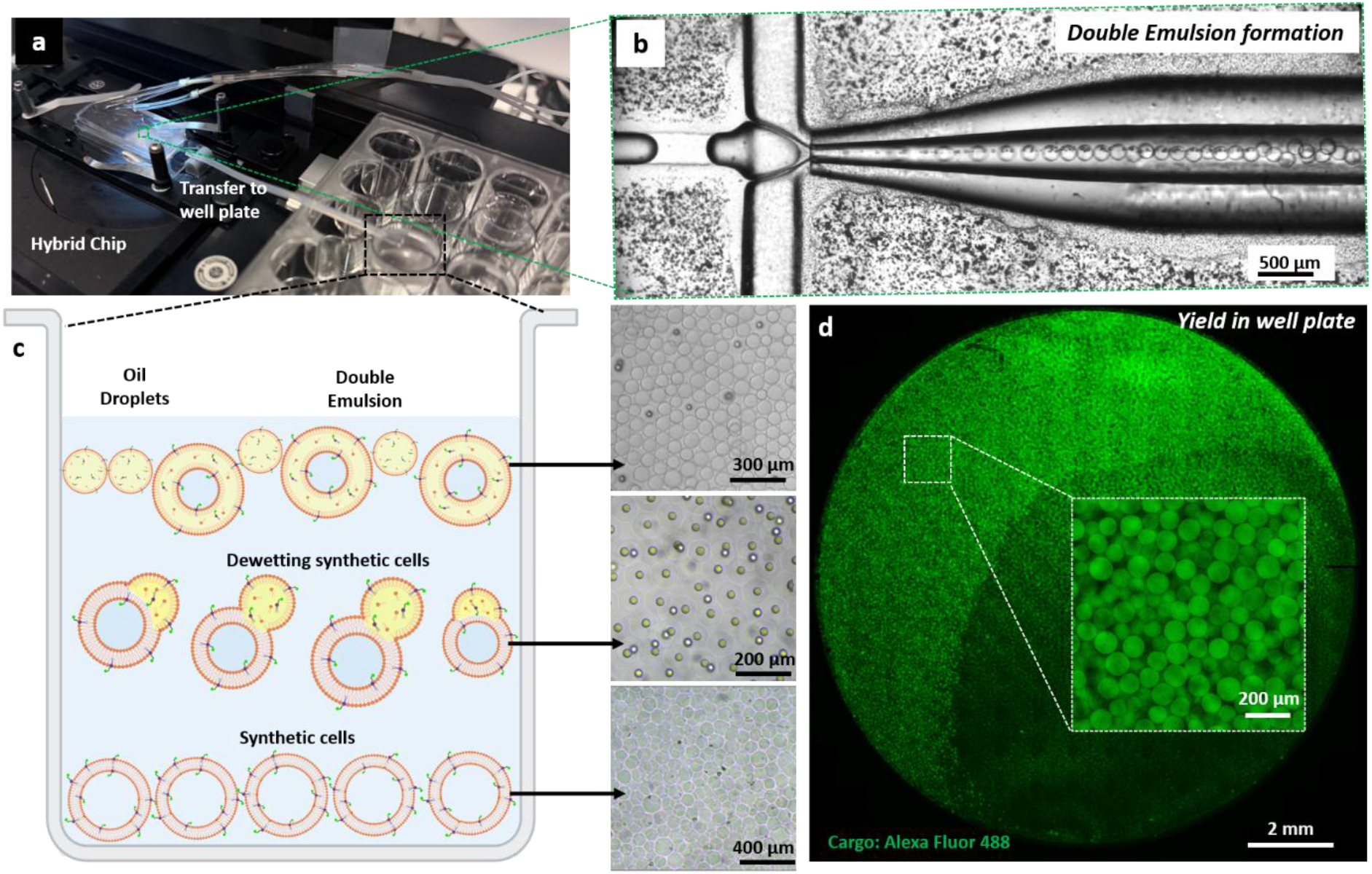
Microfluidic platform for generation of synthetic cells using Double Emulsion template method. a) Hybrid chip setup for double emulsion production. b) Brightfield image depicting the double dripping mode of double emulsion formation. c) Schematics of the events during solvent removal by dewetting of double emulsions to obtain synthetic cells. d) Confocal microscopy image at the bottom of well plate showing high yield of synthetic cells. The inset image is magnified view of a representative tile highlighting the synthetic cell yield.

**Figure 2:**
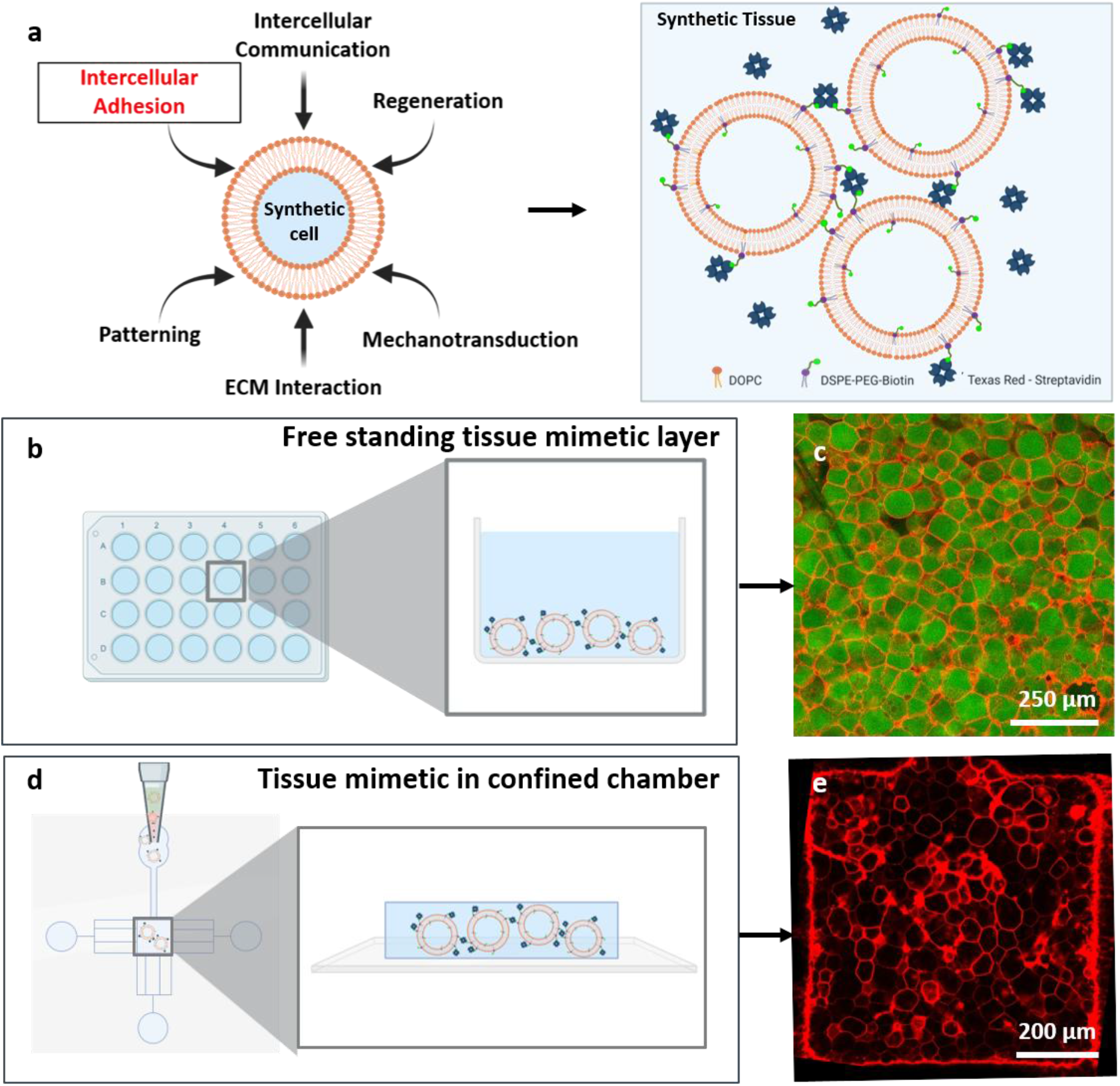
Development of synthetic tissue models. a) Functions introduced to synthetic cells to obtain a functional synthetic tissue. Schematics showing the Biotin-Streptavidin conjugation adopted in the models for intercellular adhesion. b,d) Schematic of the freestanding and confined tissue mimetic configuration respectively. c,e) Confocal micrographs of the freestanding and confined tissue mimetic configuration respectively.

### 2.2. Minimal synthetic tissue models

Synthetic cells were assembled into tissue mimic structures or minimal synthetic tissue models in two different configurations in our research. One is a freestanding monolayer in a well plate and the other is an assembly confined in a microfluidic chamber. Comparison between these two systems distinguish tissue intrinsic behaviour from the environmental induced response. In the freestanding configuration, the addition of streptavidin to sedimented biotinylated synthetic cells introduced intercellular adhesion and resulted in the formation of tissue mimetic monolayer after overnight incubation at 4°C. Quantitative characterisation of the packing was done using the Co-, ordination number along with morphological descriptors circularity and eccentricity. The deformability resultant of the intercellular adhesion is discussed in section 2.3

In the microfluidic chamber, biotinylated synthetic cells functionalised with streptavidin accumulated. The proximity/compaction between the synthetic cells resulted in intercellular adhesion. The restricted geometry provided additional mechanical constraints that contributes to the deformation of the synthetic cells. The confined tissue mimetic structure served as a platform to investigate transport process through densely packed assemblies. Fluorescent dye tracer was introduced and monitored in the chamber real time. The platform was utilized to study impact of confinement and adhesion on the transport in packed assemblies. The comparison between freestanding and confined tissue mimetic structures revealed altered transport pathway in the chamber.

### 2.3. Static packing properties of the tissue models/ Morphological characterization of the tissue models

Individual cells adapt their morphology in response to environmental cues and mechanical perturbations. Cell morphology is not only important for disease diagnosis and prognosis also influences emergent tissue behavior. Changes in cell shape influences cell-cell interactions, packing architecture, and force transmission, all of which contribute to emergent tissue properties. Quantifying cell shape provides a framework to link individual cellular response to structural and functional properties of tissues.^16,17^

To characterize synthetic cell morphology, scalar descriptors such as circularity and eccentricity were utilized. These parameters assist in translation of microscopic images to quantitative data that assesses cell response to environmental stimuli, cell-cell interactions and mechanical constraints.^18^ Circularity and Eccentricity were computed using the formula below:

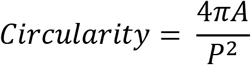

where A is the area of the synthetic cell and P is the perimeter of the cell contour^19^

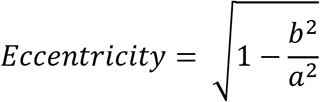

where a and b are lengths of the major and minor axes of the fitted ellipse.^20^

Circularity is 1 for a perfect circle and defined as the degree to which a shape approaches a perfect circle. Higher the circularity value, the more isotropic the shape, as regular polygons approach a circular geometry with an increasing number of edges. Eccentricity is generally used to describe nuclear morphology but can be used to assess cell polarity and elongation.^21^ Eccentricity is 1 for an ellipse and indicates the deviation from circular shape and can be used to compute anisotropic deformation or abnormal elongation resulting from mechanical stresses or confinement.

In the freestanding tissue mimetic, there was a significant difference in median of adhered and non-adhered samples (cf. Fig 3b and 3d). As expected, the non-adhered sample retained spherical morphology, exhibiting median circularity of 0.86 and median eccentricity of 0.35. On adhesion the cells deviated from spherical to polygons shapes. This transition was reflected in reduction of median circularity to 0.77 and increase in median eccentricity to 0.64. In the confined chamber, a similar trend was observed. Median circularity 0.82 and median eccentricity 0.46 for non – adhered, while for adhered tissue mimetic in chamber the median circularity was 0.77 and the median eccentricity was 0.66. These results demonstrate intercellular adhesion promotes deformation irrespective of the tissue mimetic configuration.

**Figure 3:**
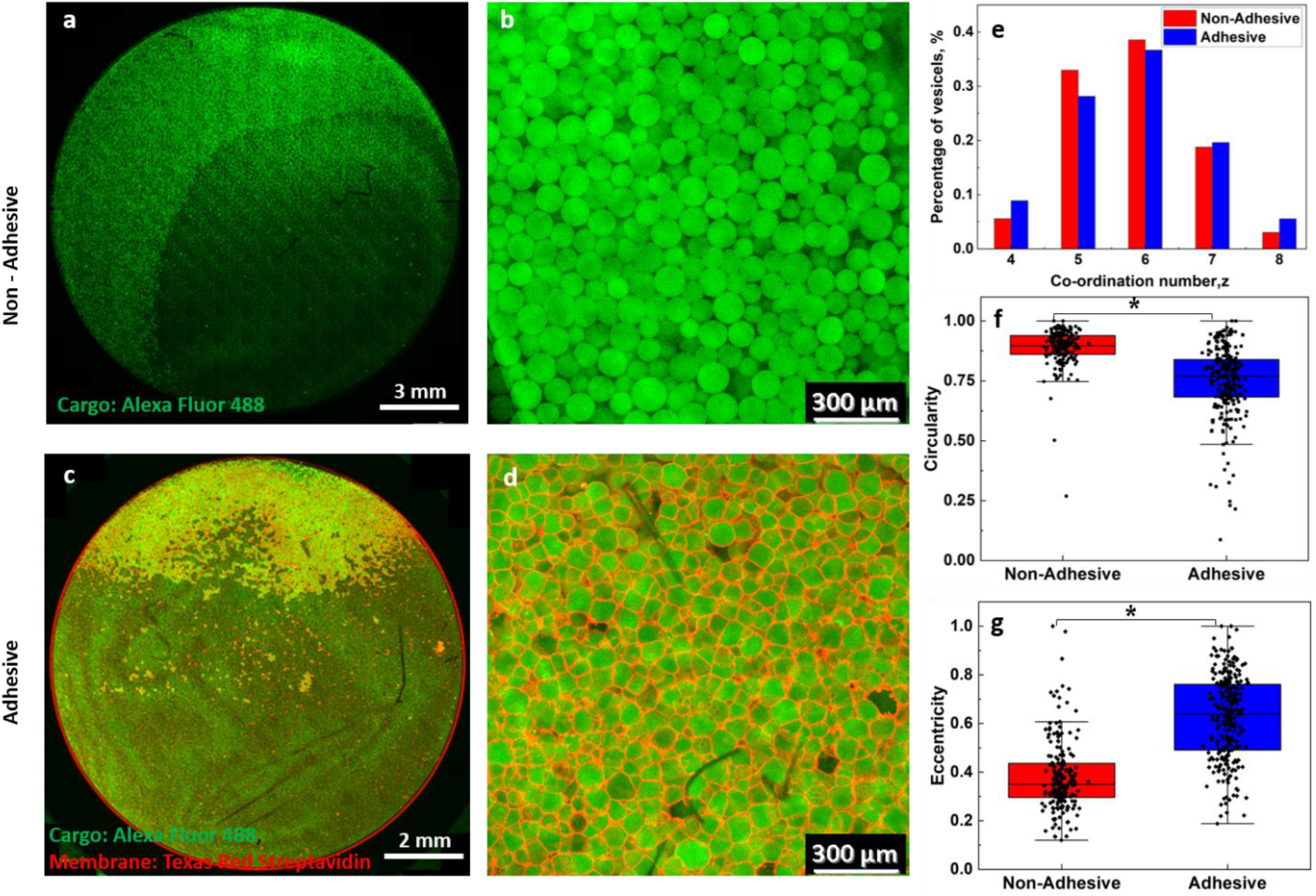
Adhesion-induced morphological changes in freestanding tissue mimetics a) Non-adhered synthetic cells in a well plate. b) Representative image of the non-adhered synthetic cells in well plate used for analysis. c) Synthetic cell-based adhered tissue mimetic in a well plate. d) Representative image of the adhered synthetic cells in well plate used for analysis. e) Co-ordination distributions. f) Circularity distributions. g) Eccentricity distributions. * p-values are calculated from Mann-Whitney test: nsp > 0.05.

Eccentricity exhibited greater variability as opposed to circularity for all samples. And circularity remained relatively clustered, suggesting eccentricity to be more sensitive. Visual inspection of the captured images supported this observation. We noticed that eccentricity effectively captured even minute deviations from isotropy, whereas circularity failed to do so.

To evaluate the effect of confinement on synthetic cell shape, the adhered populations from freestanding and confined tissue mimetics were compared. Interestingly, no statistically significant differences were detected for both the parameters (See Fig 5b and 5c). We hypothesized higher deformation in confined tissue mimetic due to the geometric restriction and flow induced elongation, but both the populations were statistically not significant. Further probing is needed to determine whether confinement effect emerges at different experimental conditions.

Local packing organization was quantified through co-ordination number (z) defined using Delaunay neighbors (See Fig 3e and Fig 4c). Although z can be calculated using geometric threshold or radial distribution function, we have used the Delaunay triangulation

**Figure 4:**
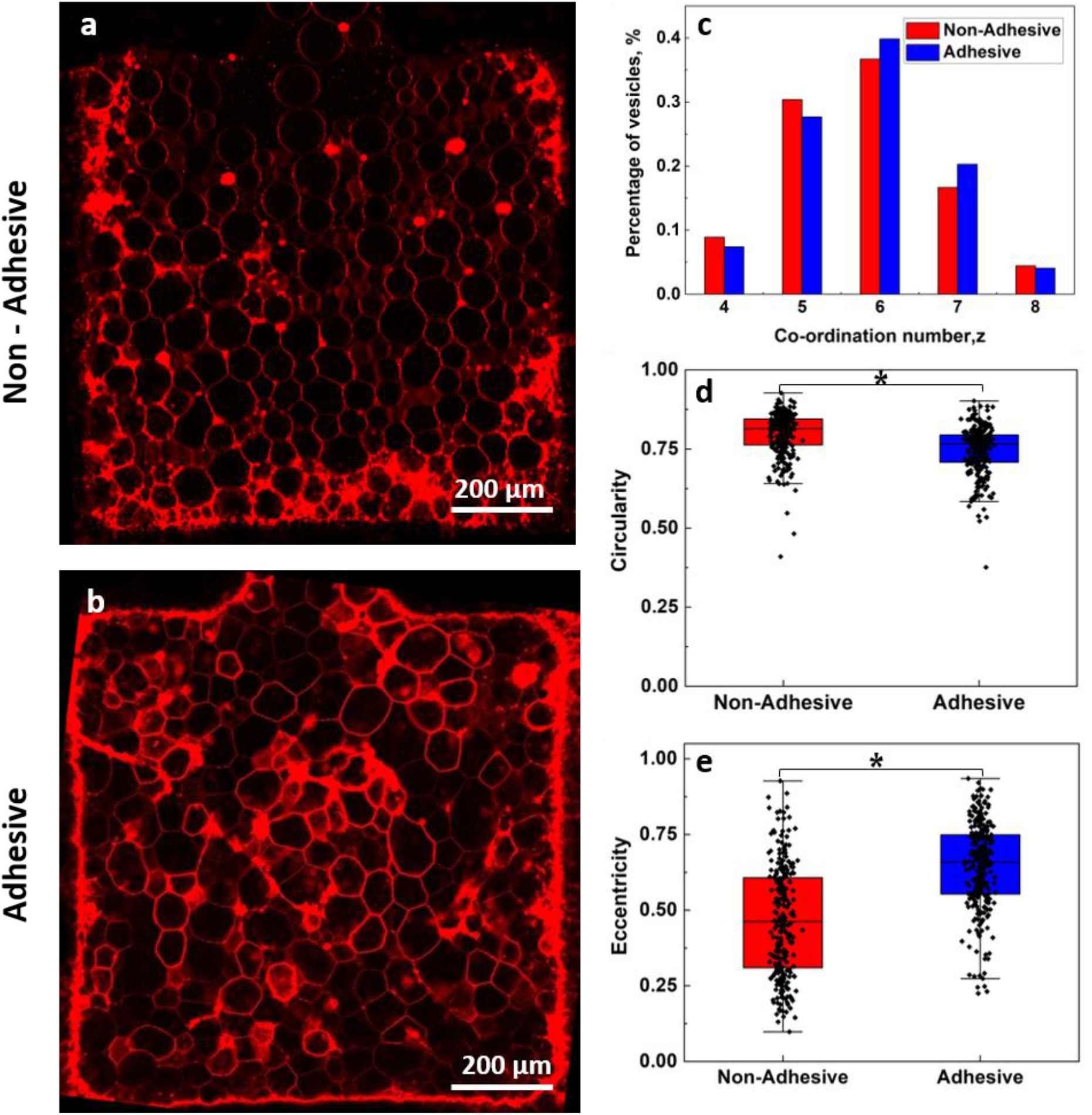
Morphological characterization of confined synthetic tissue mimetics. a) Non-adhered synthetic cells packed in a microfluidic chamber. Membrane labelling: Liss Rhod PE. b) Synthetic cell-based adhered tissue mimetic in a microfluidic chamber. Membrane labelling: Texas Red Streptavidin. c) Co-ordination numbers distributions. d) Circularity distributions. e) Eccentricity distributions. * p-values are calculated from Mann-Whitney test test: nsp > 0.05.

**Figure 5:**
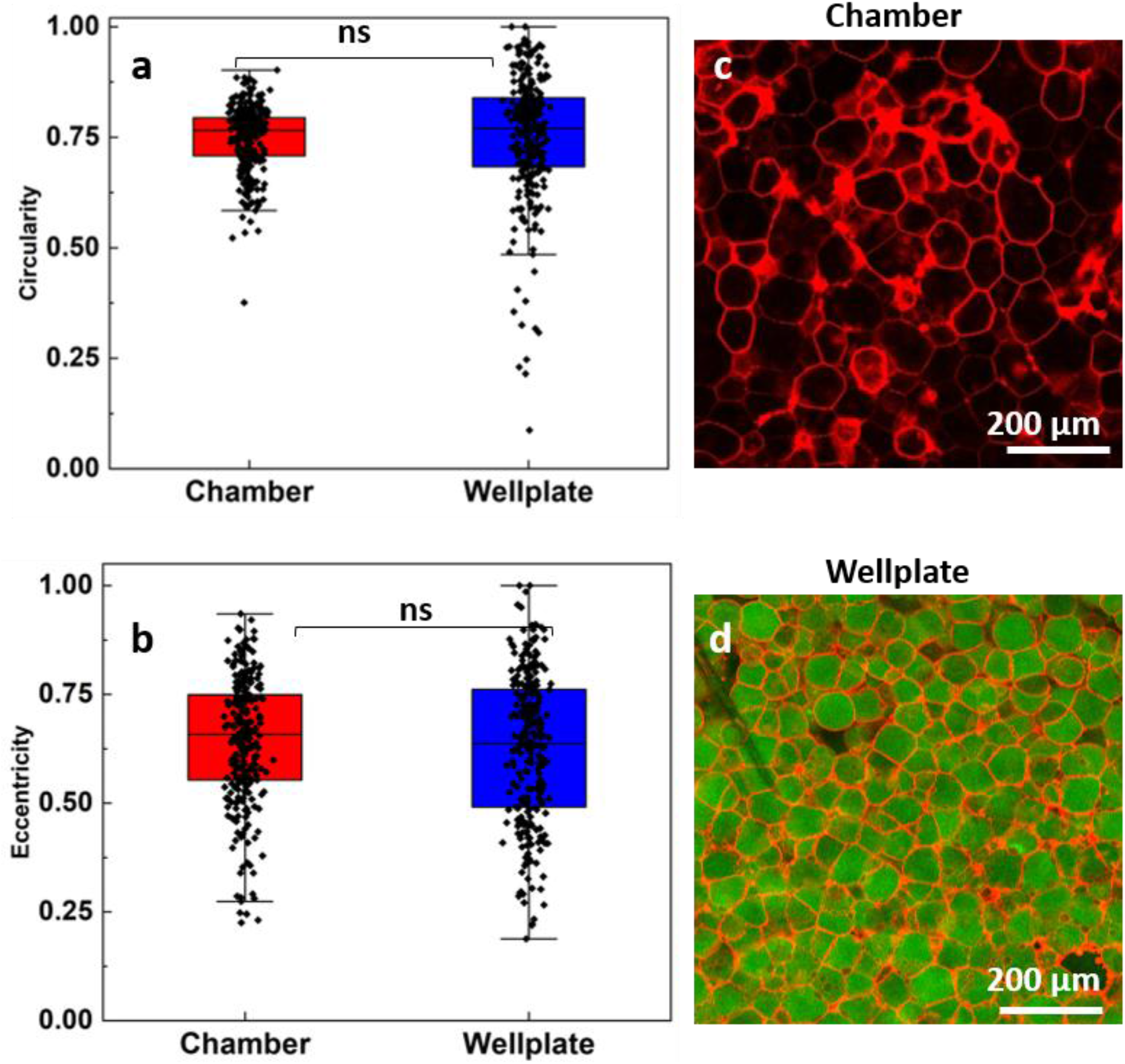
Comparison of cell morphology between freestanding and confined tissue mimetics a) Comparison of circularity distributions between the two tissue-mimetic configurations. b) Comparison of eccentricity distributions between the two tissue-mimetic configurations. c,d) Representative images of adhered tissue mimetics formed in the freestanding and confined configurations respectively.

**Figure 6:**
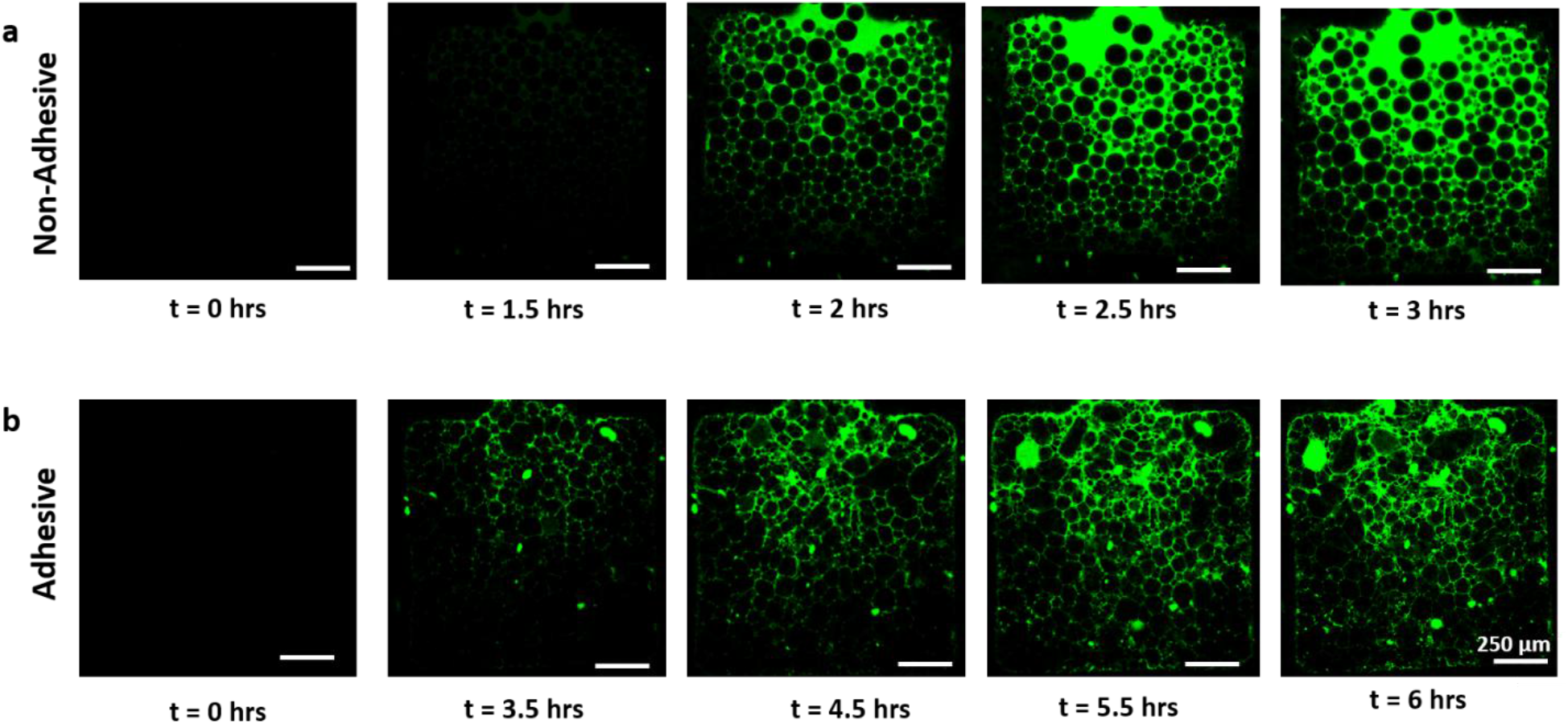
Fluorescent dye assay for transport in confined tissue mimetic. a) Advection dominated transport in non-adhered synthetic cell packing. b) Slow tortuous transport in adhesive confined tissue mimetic. Alexa Fluor 488 was used in the assay.

### 2.4. Transport properties in tissue mimetic model

We conducted the dye diffusion assay as proof-of-concept demonstration for nutrient transport or drug penetration in the developed tissue mimetic model. Fluorescent dye was introduced in the pipette tip, enabled it to enter the chamber of packed synthetic cells by the hydrostatic pressure head of the liquid. The dye transport in non-adhered sample was characterized by an advancing front consistent with advection-diffusion transport in an open packing structure.

In contrast, the adhered tissue mimetic had substantially slower transport and occurred through the interstices between packed synthetic cells. Intercellular adhesion reduced the pore size and increased the tortuosity of transport paths. Notably, the tissue mimetic remained stable throughout the experiment despite the flow. For both samples there was minimal escape and no significant bursting of synthetic cells. The confined chamber configuration packing has demonstrated the transport resistance and mechanical integrity required to model bio analyte transport and interstitial fluid (ISF) studies in packed tissues.

## 3. Conclusion

A robust microfluidic method was optimized to achieve some of the highest throughputs reported in microfluidics literature. utilized these building blocks to construct two complementary minimal tissue-mimetic systems: a freestanding assembly and a confined microfluidic tissue mimic. These platforms enabled quantitative investigation of the relationship between microscale cellular interactions and emergent tissue-scale properties. Morphometric analysis demonstrated that adhesion-mediated packing induced significant cellular deformation, resulting in distinct changes in circularity and eccentricity. Comparison of the freestanding and confined tissue mimetics revealed that adhesive interactions were the primary cause for cell deformation, while geometric confinement had a comparatively limited effect under the conditions investigated. The dye transport assay showed that intercellular adhesion in confined tissues altered transport behavior by reducing effective pore size and increasing pathway tortuosity. Thereby generating transport resistance while maintaining structural integrity under flow. By reducing biological complexity to a controllable synthetic system, we hope these platforms enable researchers to investigate the fundamental insights into collective tissue behaviors and establish design principles of functional synthetic tissues.

## 4. Methods and Materials

### Materials

Polyethylene glycol (PEG,powder/flakes), MW = 6000 gmol^−1^ was obtained from Alfa Aesar. Poloxamer (P188, 10% solution), Alexa Fluor 488 and Molecular Probes™ Streptavidin, Texas Red™ conjugate procured from Thermo Fisher. 2-[methoxy(polyethyleneoxy)propyl] trimethoxysilane was obtained from Gelest. DOPC (1,2-di-(9Z-octadecenoyl)-sn-glycero-3-phosphocholine), Liss Rhod PE (1,2-dioleoyl-sn-glycero-3-phosphoethanolamine-N-(lissamine rhodamine B sulfonyl(ammonium salt))) and DSPE-PEG (2000) Biotin (1,2-distearoyl-sn-glycero-3-phosphoethanolamine-N-[biotinyl(polyethyleneglycol)-2000] (ammonium salt)) purchased from Avanti Polar Lipids. Polydimethylsiloxane (PDMS) and curing agent (Sylgard 184™ Silicone Elastomer Kit) procured from Dow Corning. Phosphate buffered saline (PBS) from EuroClone. Glycerol, 1-Hexanol and paraffin oil were purchased from Sigma Aldrich.

### Hybrid PDMS/Glass Capillary chip and Synthetic cell Packing chip fabrication

The DE generation device used in this study is a hybrid PDMS/glass capillary system enabling prolonged generation. This device also allows for simple capillary replacement, offering a robust platform for generating double emulsions with high throughput. The fabrication of the device and capillary silane treatment for long term use has been described in detail in our earlier study.^14^ The liposome packing chip mold was fabricated via stereolithographic (SLA) 3D printing (Form 3+, Formlabs; Grey Resin; 25 μm resolution). The replicas were produced by casting a mixture of PDMS elastomer and curing agent (10:1, w/w) onto fabricated molds, followed by degassing and curing at 80 °C for 2 h. The PDMS replicas were then bonded to glass slides via oxygen plasma treatment (Colibri, Gambetti). The chips were ready to use immediately. To retain hydrophilicity, no heating post bonding was done.

### Synthetic cells generation and incubation

DE generation requires three phases. The lipid phase consisted of containing 8 mg mL^−1^ phospholipids (DOPC, DSPE-PEG Biotin 3 mol%) and 5 µg mL^−1^ Liss Rhod PE (in case of non-adhered sample) in a 60:40 (vol%) mixture of hexanol and paraffin oil. The solutions preparation protocol, inner and outer aqueous phase compositions and DE production steps can be referred to in our previous article.^15^ Streptavidin stock solution is prepared in PBS (1 mg/mL), aliquoted and stored at −20° C. Streptavidin is thawed on ice and diluted to 500 µg/mL in outer aqueous phase for the well plate tissue mimetic. This is further diluted to 40 µg/mL in the well plate containing synthetic cells and incubated overnight at 4° C to obtain the freestanding tissue configuration. For confined configuration the pipette is filled with 150 µL 40 µg/mL streptavidin solution, then 150 µL of synthetic cells suspension is added. The biotinylated cells are coated with streptavidin during sedimentation and slow into the chamber subsequently. As these synthetic cells are packed and compressed in the chamber adhesion begins and the bond is strengthened during the overnight incubation at 4° C.

### Imaging and Image processing

Images and time lapse images for the dye transport assay for analysis were obtained from Leica Falcon SP8 confocal microscope. These images were segmented using Cellpose algorithm.^22^ The segmented image is then analyzed, and the shape descriptors are calculated.

## Acknowledgements

This work was funded by the Academy of Finland (later named as Research Council of Finland), NanoEngineered Self-Assembling Vesicle Production Line (NESAV) project, grant no. 342448, and The University of Oulu & The Research Council of Finland, Profi 5, Decision no. 326291. C.E was supported in part by VTT-Technical research center of Finland Ltd, iBEX Innovation Program (ArtOfCell 2021 project). We acknowledge the assistance by Amir Mohamadsharifi Renani and Veli-Pekka Ronkainen on improvement of microfluidic device and imaging, respectively. We also acknowledge support from Biocenter Oulu Light Microscopy Core Facility, University of Oulu and Biocenter Finland. Schematic illustrations were created in https://BioRender.com. Graph plots were created using Origin 2025b.

## Use of Generative AI

Google Gemini and Google AI studio were used for code development. The authors have validated the code manually

## Data Availability Statement

Data set and code used for analyses will be shared on request.

